# abPOA: an SIMD-based C library for fast partial order alignment using adaptive band

**DOI:** 10.1101/2020.05.07.083196

**Authors:** Yan Gao, Yongzhuang Liu, Yanmei Ma, Bo Liu, Yadong Wang, Yi Xing

**Affiliations:** Center for Bioinformatics, Harbin Institute of Technology, Harbin, Heilongjiang 150001, China; Center for Computational and Genomic Medicine, Children’s Hospital of Philadelphia, Philadelphia, PA 19104, USA; Department of Pathology and Laboratory Medicine, University of Pennsylvania, Philadelphia, PA 19104, USA

## Abstract

**Summary:** Partial order alignment, which aligns a sequence to a directed acyclic graph, is now frequently used as a key component in long-read error correction and assembly. We present abPOA (**a**daptive **b**anded **P**artial **O**rder **A**lignment), a Single Instruction Multiple Data (SIMD) based C library for fast partial order alignment using adaptive banded dynamic programming. It can work as a stand-alone multiple sequence alignment and consensus calling tool or be easily integrated into any long-read error correction and assembly workflow. Compared to a state-of-the-art tool (SPOA), abPOA is up to 15 times faster with a comparable alignment accuracy.

**Availability and implementation:** abPOA is implemented in C. A stand-alone tool and a C/Python software interface are freely available at https://github.com/yangao07/abPOA.

## 1 Introduction

Partial order alignment (POA) was first introduced by Lee *et al.* (2002) to solve the multiple sequence alignment (MSA) problem. In POA, MSA is represented as a directed acyclic graph (DAG) and sequences are iteratively aligned to the DAG through dynamic programming (DP). Multiple consensus sequences are then generated by applying the heaviest bundling algorithm to the alignment graph (Lee, 2003). Recently, with the advent and growing popularity of long-read sequencing technologies on the Pacific Biosciences (PacBio) and Oxford Nanopore Technologies (ONT) platforms, there is a renewed appreciation and interest of POA, and this algorithm is now broadly used for error correction and assembly of error-prone long reads (Loman *et al.*, 2015; Vaser *et al.*, 2017; Volden *et al.*, 2018; Gao *et al.*, 2019; Ruan and Li, 2019).

Although POA is much faster than classical MSA algorithms (Lassmann and Sonnhammer, 2002), the large datasets generated by state-of-the-art long-read sequencing platforms pose a major challenge. To address this challenge, Single Instruction Multiple Data (SIMD) implementation was used to accelerate the original POA algorithm (Vaser *et al.*, 2017). This SIMD version of POA, SPOA, takes advantage of the wider SIMD registers in modern processors that process multiple elements in parallel. SIMD vectors are used to store scores of multiple consecutive cells in each row of the DP matrix and processed using SIMD instructions, with parallel updating for all scores stored in each vector.

In addition to the SIMD parallelization, another acceleration strategy, “banded DP”, is also widely used in sequence-to-sequence alignment tools (Chao *et al.*, 1992). Specifically, in each row or column of the DP matrix, only cells inside a specific “band” need to be filled out. Recently, a graph version of this strategy was explored in GraphAligner (Rautiainen and Marschall, 2019), in which a band for each row of the DP matrix is dynamically defined based on the minimum score cell in that row. Other cells are considered to be inside the band only if the score is within a certain distance of that minimum score. Although GraphAligner is faster than other graph alignment tools, its strategy may not be suitable for MSA and consensus calling of long reads with high error rates. This is due to GraphAligner’s using of edit distance instead of general scoring function as the alignment metric, which may lead to incorrect alignments in regions with a large number of insertion or deletion errors that are common in PacBio and ONT sequencing data.

In this work, we have developed abPOA, an extended version of POA that performs adaptive banded DP with an SIMD implementation. abPOA supports flexible scoring schemes. It can work as a stand-alone MSA and consensus calling tool, or be easily integrated into any long-read error correction and assembly workflow.

## 2 Methods

abPOA adopts the same SIMD parallelization strategy as in SPOA, where the SIMD vectors are placed parallel to the linear sequence. In the DP matrix, each row corresponds to one node in the alignment graph and each column corresponds to one base of the linear sequence being aligned (Fig. 1A and Supplementary Fig. 1). Instead of filling out the entire DP matrix, abPOA adaptively defines a band for each row based on the scores in predecessor rows and the lengths of potential outgoing paths in the alignment graph, and only scores of cells inside this band are calculated.

**Figure 1:**
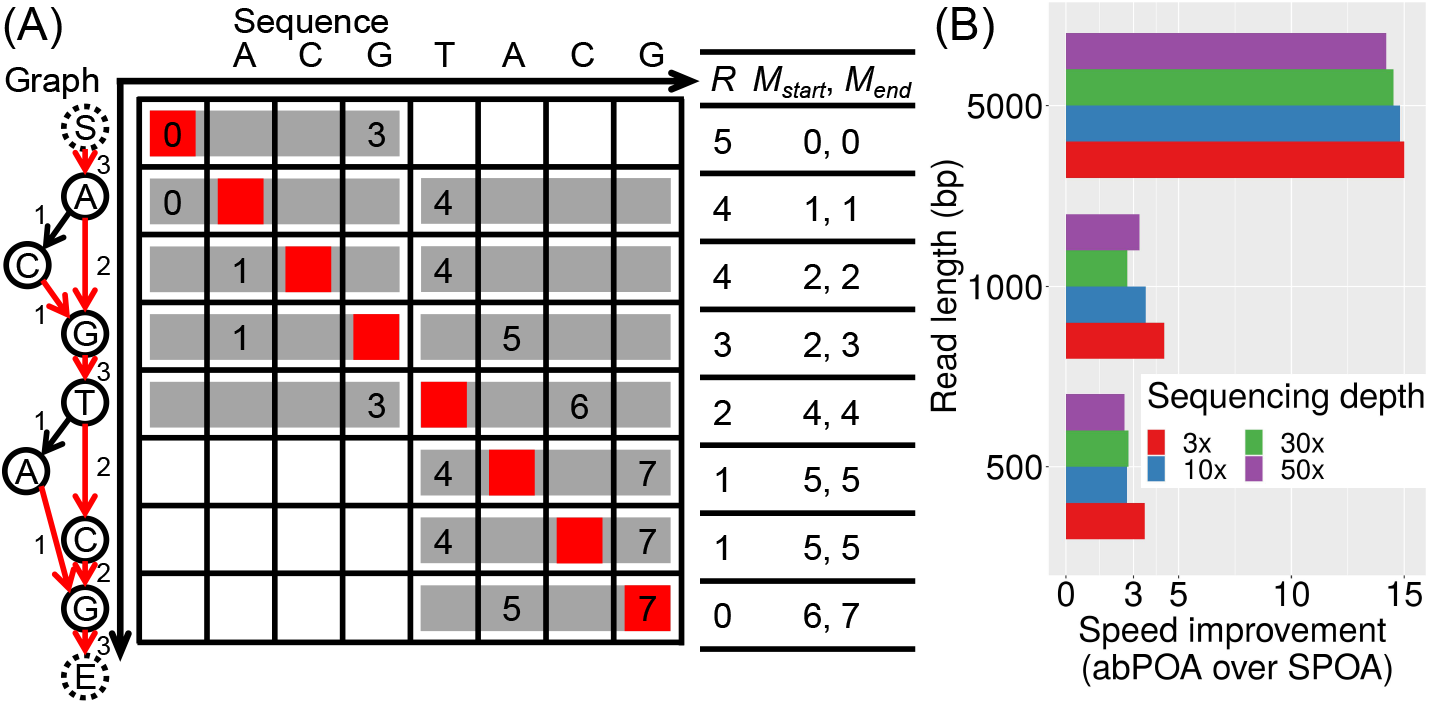
(A) Illustration of the SIMD parallelization and adaptive banded DP applied in abPOA. In the alignment graph, ‘S’ is the start node and ‘E’ is the end node. The supporting read count for each edge of the alignment graph is shown next to the edge. For each node, the heaviest outgoing edge is marked in red. Inside the DP matrix, the two numbers in each row are the base-level boundary of the band in that row. Grey blocks represent SIMD vectors that contain bases inside the band. In this example, each SIMD vector is composed of four consecutive DP scores in each row. The red block represents the maximum score cell of each row. The values of *R*, *M*_*start*_, and *M*_*end*_ are shown on the right side of the matrix. Besides, *L* is 7 and *w* is 1 in this example. (B) The speed improvement of abPOA with adaptive banding over SPOA on 12 sequence sets simulated by NanoSim.

In more detail, for each node in the alignment graph, abPOA first computes the length of the outgoing path with the largest number of supporting reads that starts from the current node to the end node. This length, *R*, is considered as the most likely number of additional bases to be included in the alignment path starting from the current node. abPOA iteratively calculates *R* for all nodes in a similar way to the heaviest bundling algorithm (Lee, 2003) (Supplementary Note). Note that *R* is calculated before each round of the sequence-to-graph alignment, based on the nodes and edge weights of the current alignment graph. Then, during the DP process, all rows of the DP matrix are sequentially processed following the partial order of the graph. As such, for each row, abPOA can collect the horizontal coordinates of maximum score cells in its predecessor rows, i.e. their positions in the linear sequence. The range (start and end positions) of possible maximum score cells in the current row, [*M*_*start*_, *M*_*end*_], can be derived as *M*_*start*_ = *P*_*left*_ + 1 and *M*_*end*_ = *P*_*right*_ + 1, where *P*_*left*_ and *P*_*right*_ are the positions of the leftmost and the rightmost maximum score cells in all predecessor rows. *M*_*start*_ and *M*_*end*_ of the first row are set as 0 since the start node has no predecessor.

With the above numbers calculated for each row, abPOA defines the start and end positions of the DP band in each row as *B*_*start*_ = max{0, min {*M*_*start*_, *L* − *R*} − *w* and *B*_*end*_ = min{*L*, max{*M*_*end*_, *L* − *R* + *w*}, where *L* is the length of the linear sequence and *w* is the number of extra bases added on both sides of the band, which is determined by two parameters *b* and *f* (default 10 and 0.01) as *b* + *f* × *L* (Fig. 1A and Supplementary Note). By taking both the scores in predecessor rows and potential outgoing alignment paths into consideration, this adaptively defined DP band is expected to fully cover the optimal alignment path, even for divergent sequences and graphs. Next, abPOA maps the base-level boundary of the adaptive band onto SIMD vectors to only process vectors that contain bases inside the band. After iteratively aligning sequences to the graph and updating the graph (Lee *et al.*, 2002), abPOA generates a consensus sequence from the final alignment graph using the heaviest bundling algorithm (Lee, 2003).

## 3 Result

We evaluated abPOA using simulated long-read datasets along with SPOA (Vaser *et al.*, 2017), which to our knowledge is the only existing tool that uses SIMD to accelerate POA. Three read lengths (300 bp, 1000 bp, and 5000 bp) and four sequencing depths (3×, 10×, 30×, and 50×) were used to simulate 12 sets of sequences each with 100 clusters of sequences to be aligned, using NanoSim (Yang *et al.*, 2017) or PBSIM (Ono *et al.*, 2012) to incorporate error profiles of ONT or PacBio respectively (Supplementary Tables 1 and 2). Each cluster of sequences was aligned by abPOA or SPOA to generate a consensus sequence, and abPOA was run twice with adaptive banded DP enabled or disabled (see details of the evaluation procedure in Supplementary Note).

abPOA is 2.6–15.0 times faster than SPOA on NanoSim simulated sequence sets when adaptive banded DP is enabled (Fig. 1B and Supplementary Table 1). As expected, the speed improvement of abPOA over SPOA becomes particularly pronounced for longer sequences, as more SIMD vectors would be skipped outside of the adaptive band leading to a more significant efficiency gain. A similar trend can be observed from the PBSIM simulation (Supplementary Table 2). To evaluate alignment accuracy, we calculated the error rate of the generated consensus sequence (Supplementary Note). abPOA yields a comparably low error rate of consensus sequences when compared to SPOA (Supplementary Tables 1 and 2). Moreover, compared to abPOA without adaptive banding, abPOA with adaptive banding significantly reduces the run time i n all simulation settings without sacrificing the alignment accuracy.

In summary, our results demonstrate that abPOA can generate high-quality consensus sequences from error-prone long reads and offer significant speed improvement over existing tools. With the significant impact of POA on third-generation long-read sequencing data analysis, we expect that abPOA will be a useful and broadly applicable tool in long-read bioinformatics workflows.

## Acknowledgements

The authors would like to thank Dr. Yuan Gao and Yadong Liu for assistance with testing abPOA and providing comments on the manuscript.

---

**Supplementary Fig. 1.**
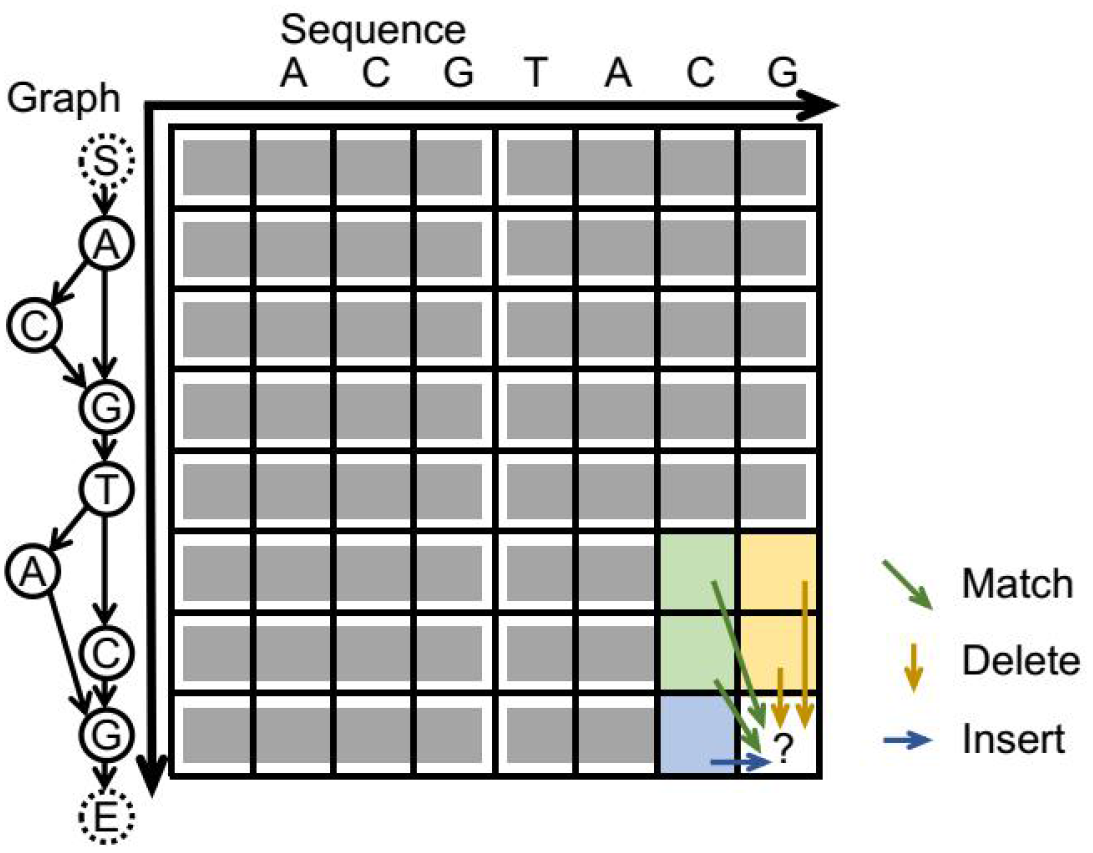
Illustration of the SIMD parallelization applied in abPOA and three types of DP operations to be processed. In the alignment graph, ‘S’ is the start node and ‘E’ is the end node. Inside the DP matrix, each row corresponds to one node in the alignment graph and each column corresponds to one base of the linear sequence being aligned to the graph. Grey blocks represent SIMD vectors each composed of four consecutive DP scores in each row. Note that abPOA can adaptively determine the number of elements to be stored in each vector based on the size of the register available in the computer processors and the lengths of the aligned sequences. abPOA processes all the vectors in a row-by-row manner following the partial order of the graph. During the DP process, for “match” and “delete” operations (diagonal and vertical moves in the DP matrix), all scores stored in each SIMD vector can be updated in parallel as they only rely on scores in the predecessor rows. For “insert” operations (horizontal moves in the DP matrix), sequential non-parallel updating of scores in the same SIMD vector is needed, as the score of each cell depends on the score of the cell on the left.

**Supplementary Table 1.** Run time^a^ (CPU sec) and consensus sequence’s error rate^b^ (%) of abPOA and SPOA on 12 sequence sets simulated by NanoSim.

| Length × depth<br>(error rate %) | SPOA | abPOA without<br>adaptive banding | abPOA with adaptive<br>banding |
| --- | --- | --- | --- |
| 500 × 3 (11.69) | 0.14<sup>a</sup> (4.68<sup>b</sup>) | 0.11 (4.60) | 0.04 (4.61) |
| 500 × 10 (11.75) | 0.65 (0.24) | 0.66 (0.27) | 0.24 (0.27) |
| 500 × 30 (11.67) | 3.41 (0.05) | 3.18 (0.05) | 1.23 (0.05) |
| 500 × 50 (11.56) | 7.18 (0.09) | 6.59 (0.07) | 2.76 (0.07) |
| 1000 × 3 (12.05) | 0.48 (4.55) | 0.45 (4.54) | 0.11 (4.54) |
| 1000 × 10 (12.23) | 2.41 (0.24) | 2.50 (0.27) | 0.68 (0.27) |
| 1000 × 30 (12.12) | 11.82 (0.05) | 12.13 (0.04) | 4.34 (0.04) |
| 1000 × 50 (12.18) | 23.17 (0.09) | 29.69 (0.07) | 7.10 (0.07) |
| 5000 × 3 (12.86) | 16.35 (4.94) | 10.48 (4.89) | 1.09 (4.89) |
| 5000 × 10 (12.93) | 98.50 (0.31) | 60.01 (0.30) | 6.65 (0.30) |
| 5000 × 30 (12.94) | 472.29 (0.07) | 286.51 (0.07) | 32.52 (0.07) |
| 5000 × 50 (12.92) | 1,013.08 (0.08) | 766.38 (0.06) | 71.30 (0.06) |

*Notes:* Run time is summed up for each sequence set over the 100 clusters of sequences to be aligned. Error rates of simulated long-read sequences and generated consensus sequences are shown inside the parentheses, as calculated based on the alignments to the original sequences using minimap2 (Li, 2018) (Supplementary Note) and averaged across each sequence set.

**Supplementary Table 2.** Run time^a^ (CPU sec) and consensus sequence’s error rate^b^ (%) of abPOA and SPOA on 12 sequence sets simulated by PBSIM.

| Length × depth<br>(error rate %) | SPOA | abPOA without<br>adaptive banding | abPOA with adaptive<br>banding |
| --- | --- | --- | --- |
| 500 × 3 (10.93) | 0.12<sup>a</sup> (8.19<sup>b</sup>) | 0.11 (8.15) | 0.04 (8.15) |
| 500 × 10 (12.55) | 0.64 (2.36) | 0.67 (2.62) | 0.25 (2.62) |
| 500 × 30 (12.59) | 3.01 (1.71) | 3.55 (1.76) | 1.37 (1.76) |
| 500 × 50 (12.56) | 6.91 (1.77) | 6.91 (1.61) | 2.96 (1.61) |
| 1000 × 3 (12.08) | 0.44 (8.33) | 0.42 (8.28) | 0.10 (8.28) |
| 1000 × 10 (12.99) | 2.38 (2.46) | 2.57 (2.73) | 0.66 (2.73) |
| 1000 × 30 (13.08) | 11.91 (1.94) | 12.95 (1.92) | 3.60 (1.93) |
| 1000 × 50 (12.86) | 26.41 (1.69) | 30.82 (1.60) | 7.27 (1.60) |
| 5000 × 3 (13.33) | 20.88 (9.67) | 10.37 (9.64) | 1.13 (9.64) |
| 5000 × 10 (13.49) | 107.73 (3.26) | 68.39 (3.36) | 6.63 (3.36) |
| 5000 × 30 (12.88) | 508.65 (1.50) | 308.72 (1.51) | 34.90 (1.51) |
| 5000 × 50 (12.84) | 1,096.99 (1.45) | 770.95 (1.42) | 75.91 (1.42) |

*Notes:* Run time is summed up for each sequence set over the 100 clusters of sequences to be aligned. Error rates of simulated long-read sequences and generated consensus sequences are shown inside the parentheses, as calculated based on the alignments to the original sequences using minimap2 (Li, 2018) (Supplementary Note) and averaged across each sequence set.

## Supplementary Note

### 1. Calculation of *R*

Before each round of the sequence-to-graph alignment, abPOA computes *R*, the length of the outgoing path with the largest number of supporting reads that starts from the current node to the end node. Note that neither the current node nor the end node is counted in the path. abPOA iteratively calculates *R* for all nodes in a similar way to the heaviest bundling algorithm in POA (Lee, 2003). Algorithm 1 gives the pseudo-code to calculate *R* through a graph traversal starting from the end node (line 5-6) to the start node (line 19-20). Here, the start node has no predecessor and the end node has no successor, and they both are auxiliary nodes that have no sequence bases (Fig. 1A and Supplementary Fig. 1). For each node, the outgoing edge with the heaviest weight and its corresponding successor node are picked out (line 15-17). Then, *R* of the current node is set as *R* of the chosen successor node increased by one (line 18). abPOA maintains a first-in-first-out queue and an array of all nodes’ out degrees to make sure every node is visited only after all of its successor nodes have been visited (line 23-24).

#### Algorithm 1. Calculate *R*

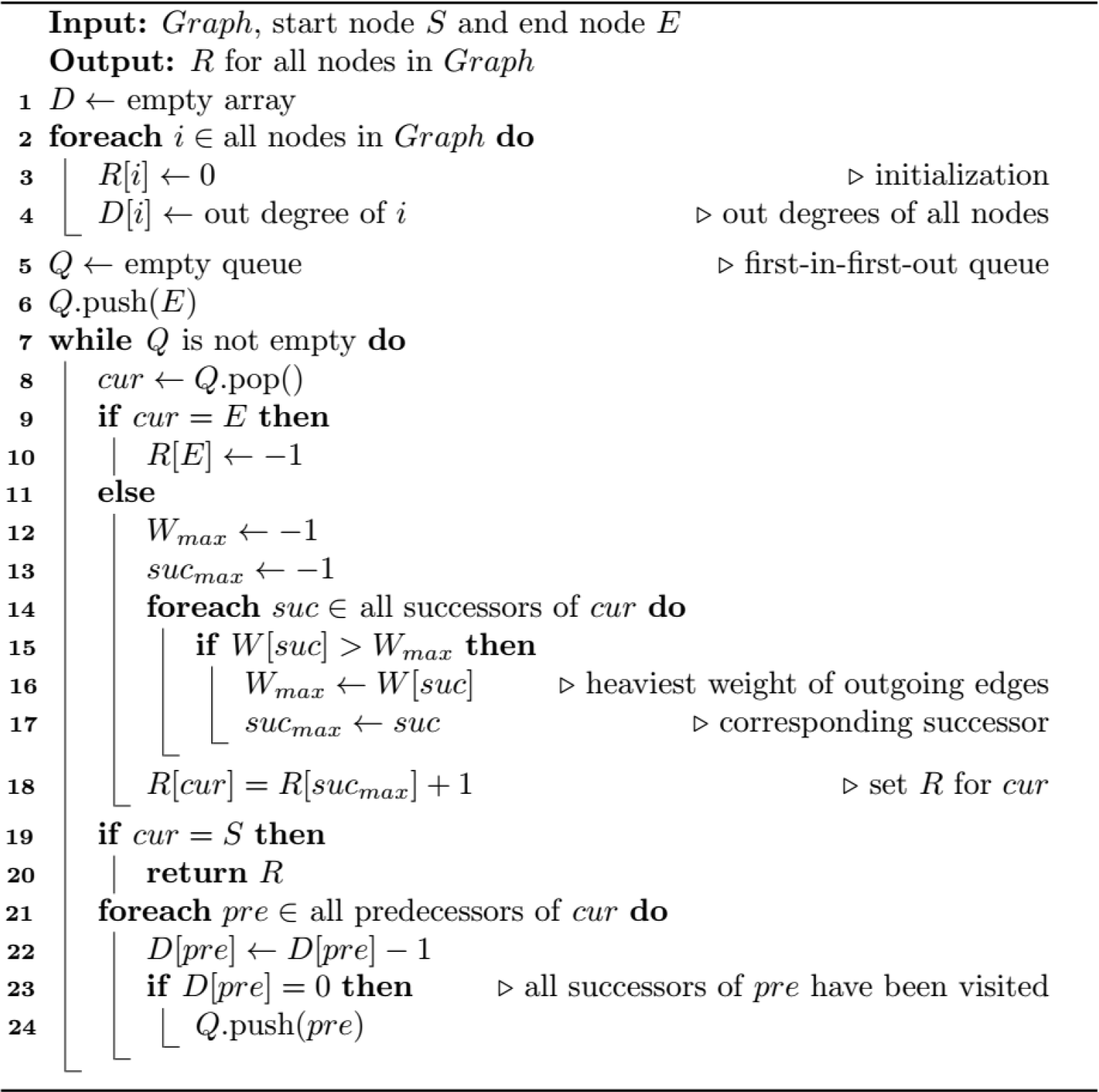

### 2. The number of extra bases added on both sides of the adaptive band

To further improve the alignment accuracy, abPOA allows the DP band in each row to be extended on both sides by *w*. Before each round of the sequence-to-graph alignment, *w*, the number of extra bases added on each side of the band is determined as *b* + *f* × *L*, where *b* and *f* are two parameters (default: 10 and 0.01), *L* is the length of the linear sequence being aligned to the graph. *w* is always rounded down to the nearest integer number.

### 3. Simulation procedure

To simulate long-read datasets with varying read lengths and sequencing depths, we first randomly extracted a sequence with a specific length from the GRCh38 human reference genome. Then NanoSim (Yang *et al.*, 2017) or PBSIM (Ono *et al.*, 2013) were used to simulate a specific number of reads from the extracted sequence. With three lengths (500 bp, 1000 bp, and 5000 bp) and four depths (3×, 10×, 30×, and 50×), a combination of 12 sequence sets were generated by each simulator. Within each simulation setting, the simulation was repeatedly run 100 times to generate 100 clusters of sequences and each cluster was aligned by abPOA or SPOA to generate a consensus sequence.

The run settings of NanoSim:

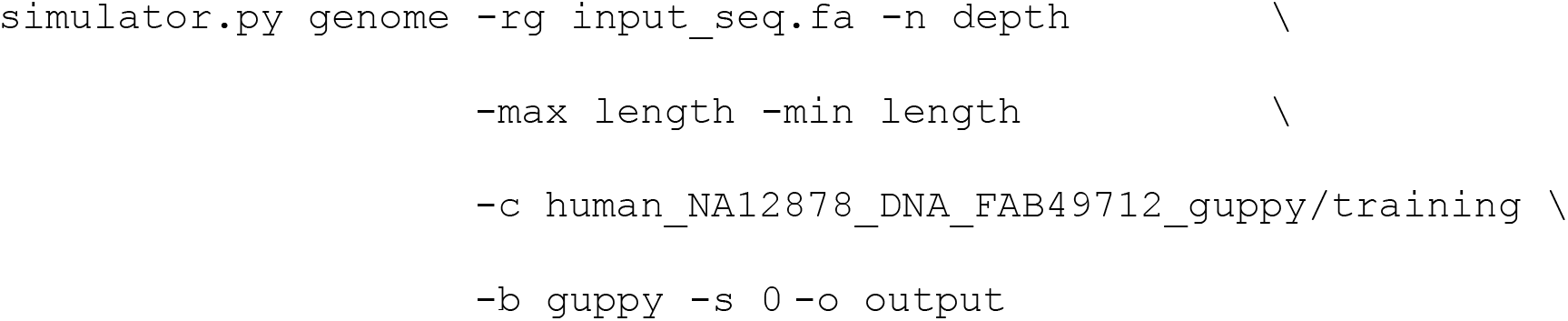

The run settings of PBSIM:

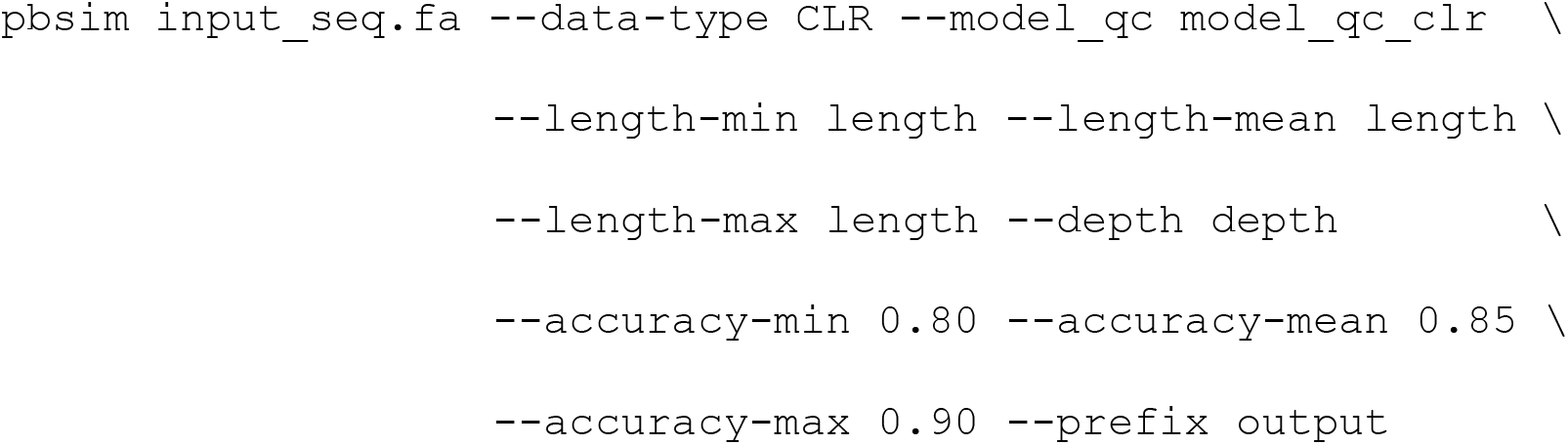

### 4. Evaluation procedure

Evaluation of abPOA and SPOA on the simulated long-read datasets was performed on a Linux system with Intel Core i5-6200U at 2.3 GHz and AVX2 instructions available. Two wrap-up programs (available at https://github.com/yangao07/abPOA) were written to make the two libraries allow each dataset that consists of 100 clusters of sequences as the input.

Both abPOA and SPOA were run in global alignment mode using a convex gap penalty scheme, i.e. two-piece gap penalty scheme. The convex penalty of a gap with length *g* is *min{O_1_+g*×*E*_*1*_, *O*_2_+*g*×*E*_*2*_}, where *O*_*1*_, *E*_*1*_, *O*_*2*_, and *E*_*2*_ are the two pairs of gap opening and gap extension penalties. They were set as *O*_*1*_=4, *E*_*1*_=2, *O*_*2*_=24, and *E*_*2*_=1 in the evaluation runs for both abPOA and SPOA.

The run settings of SPOA:

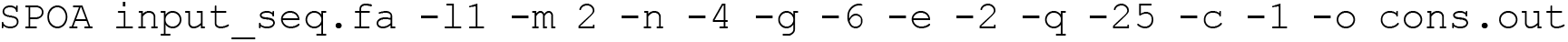

The run settings of abPOA without adaptive banding:

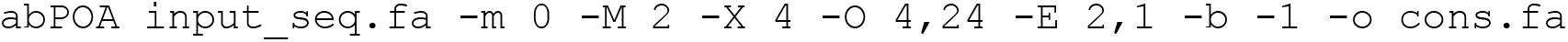

The run settings of abPOA with adaptive banding:

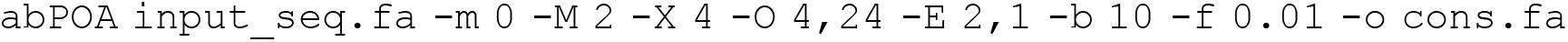

To evaluate the alignment accuracy of abPOA and SPOA, we calculated the error rate of the generated consensus sequence. We aligned each consensus sequence to the original sequence using minimap2 (Li, 2018) with default settings. The error rate is calculated as the total number of mismatches, insertions, and deletions in the alignment divided by the length of the consensus sequence. We also computed the error rate of raw simulated sequences in the same way. The averaged error rates of raw simulated sequences and generated consensus sequences across each sequence set are shown in Supplementary Tables 1 and 2.

